# The role of m6A methyltransferase WTAP in inflammatory response of atherosclerosis through NF-κB/NLRP3 mediated pyroptosis

**DOI:** 10.1101/2024.01.12.575466

**Authors:** Bing Hu, Mei He, Yanhua Sha, Fengxia Guo

## Abstract

**Background:** Pyroptosis is a new form of pro-inflammatory programmed cell death that has been linked to the development of atherosclerosis (AS). However, its exact mechanisms are not known. N6-methyladenosine (m6A) methylation is the commonest and most abundant epigenetic modification of eukaryotic mRNAs. m6A methylation modulates pathological and physiological processes involved in cardiovascular diseases. However, the exact mechanism by which it regulates inflammation in AS is unclear.

**Methods:** In this study, the level of m6A and WTAP in CHD was explored. To determine the effect of WTAP on the release of pyrolysis-related proteins and pro-inflammatory cytokines, the expression of WTAP in lipopolysaccharide (LPS)-treated endothelial cells was silenced. Pyroptosis-related proteins and pro-inflammatory cytokines were quantified in the presence of NLRP3 shRNA (shNLRP3) and NF-kB shRNA (shNF-kB(p50)). The interaction of NF-kB and NLRP3 was examined through immunoprecipitation (CO-IP), immunofluorescence (IF) and GST-pull down assays.

**Result:** It was observed that lipopolysaccharide (LPS) induced Nod-like receptor protein 3 (NLRP3)-mediated pyroptosis and inflammation, both of which were abolished through the knockdown of WTAP. Interestingly, our results indicated that WTAP enhanced the function of nuclear factor κB (NF-κB) p50 (an NF-κB subunit) and that p50 could interact with NLRP3 in endothelial cells.

**Conclusion:** In conclusion, these results suggested that WTAP in the formation of pyroptosis and inflammation in endothelial cells exposed to LPS stress by activating the NF-κB/NLRP3 signaling pathway. These findings demonstrate the mechanism of WTAP regulation during the progression of AS.

## Introduction

Current research suggests that inflammation plays an important role in the pathogenesis of atherosclerosis (AS) ^[1]^. Furthermore, the role of inflammation in the progression of plaque formation in AS has been confirmed ^[2]^. Inflammation-induced endothelial dysfunction, on the other hand, initiates and leads to the development of AS ^[3]^. Therefore, reducing inflammation in endothelial cells (ECs) may have beneficial effects for the prevention of AS progression.

Pyroptosis, also called inflammatory cell death, is a type of coordinated cellular necrosis wherein cells swell till the rupturing of their cell membranes, releasing cellular contents and activating pathways that cause severe inflammatory effects. Pyroptosis also results in the development of various cardiovascular diseases. Nicotine induces AS via activating the ROS-NLRP3 pathway, thus causing EC pyroptosis ^[4]^ and the caspase inhibitor VX-765 has been found to alleviate AS in ApoE^-/-^mice via pyroptosis modulation ^[5]^. Estrogen suppresses AS by activating estrogen receptor alpha-regulated autophagy, which reduces EC pyroptosis ^[6]^. Therefore, research involving therapeutic targets to alleviate pyroptosis is vital for the potential treatment and prevention of AS. The exact mechanism of pyroptosis in AS, however, is yet unknown.

N6-methyladenosine (m6A) methylation is the most widespread and abundant epigenetic alteration of all eukaryotic mRNAs, and is regulated by m6A methyltransferases, demethylases, and m6A-binding proteins. This methylation affects the transcription, cleavage, translation, and degradation of target mRNAs ^[7]^. Wilms tumor 1-associated protein (WTAP) is an essential component of several key m6A methyltransferases, which stabilizes METTL14 and methyltransferase-like (METTL)3^[8]^. The total panax notoginseng saponins (TPNS) was reported to modulate the WTAP/p16 signal axis through m6A modification, thereby inhibiting the balloon catheter injury-induced proliferation, migration and intimal hyperplasia of VSMCs in rat carotid arteries. This suggested that m6A modification-mediated regulation of gene expression may be a potential target for arterial restenosis ^[9]^. In some studies, the WTAP mRNA expression was found to be positively linked to pro-inflammatory cytokines and NLRP3 inflammasome components ^[10]^. Altogether, these studies suggest that WTAP may regulate atherosclerotic process by modulating pyroptosis, but the exact mechanism remain to be clarified.

In this study, we aim to investigate the exact mechanism of thermoprotein deposition in AS. m6A are important in the etiology of AS. In this study, we found that WTAP significantly enhanced pyroptosis and inflammation induced by LPS. Mechanistically, WTAP increased the NF-κB p50 activity, and p50 interacted directly with NLRP3 in endothelial cells. In summary, this study demonstrates that WTAP is a novel NF-κB-related m6A in endothelial cells that may participate in AS via NF-κB/NLRP3 signaling and may be a prospective therapeutic target for the prevention of AS.

## Materials and Methods

### Animals

All investigations conformed to the Guide for the Care and Use of Laboratory Animals published by the US National Institutes of Health (NIH Publication No. 85-23, revised 1996) and were approved by the Animal Experimental Committee of Henan Provincial People’s Hospital. Male C57BL/6 mice (6 weeks of age, 20 g) and ApoE^-/-^mice with a C57BL/6 background were obtained from the Laboratory Animal Center of Peking University (Beijing, China). To ascertain the effect of WTAP on atherosclerosis, the male (6 weeks of age) ApoE^-/-^ mice were randomized into three groups of 42 mice respectively (ND, HFD and AAV-si-WTAP-treated groups). The high-fat diet mice were injected with adenovirus si-RNA-WTAP (AAV-si-FTO) through the tail vein. Mice (6 weeks of age) were fed a high-fat diet (21% protein, 24% carbohydrate, and 55% fat) for 16 weeks. At week 16, mice were anesthetized with inhalation of 2% sodium valproate. The mice were euthanized by cervical dislocation, and tissues were collected for further analyses.

### Cell culture

The Human umbilical vein endothelial cells (Endothelial cells; ATCC CRL-1730) were procured through a local supplier from American Type Culture Collection (Manassas, VA, USA). The cells were then seeded in Dulbecco modified eagle’s medium (DMEM) containing fetal bovine serum (FBS, 10%) in 5% CO_2_ and 95% atmosphere at 37 °C. The Cells were grown in 60-mm dishes or 6-well or 12-well plates and were allowed to attain 70–80% confluency before their further use.

### Western blotting and immunoprecipitation assays

12.5% SDS-PAGE was used for separation of protein lysates and the blotted membranes were incubated with primary antibodies (1:800-dilution) for 12 hours followed by incubation with secondary antibodies for 2 hours at room temperature. Chemiluminescence was then used to detect the proteins of interest using the ECL Plus Western Blot Detection System (Amersham Biosciences, Foster City, USA)^[11]^. Immunoprecipitation was performed following the previously published protocol ^[12]^. Lysates from Endothelial cells were precleared with a mixture of 1 mg protein A/G PLUS-agarose beads and control IgG, followed by their overnight incubation at 4 °C with antibodies (anti-NLRP3) and protein A/G PLUS-agarose beads (25 mL). Using the SDS-PAGE sample buffer, the obtained immunoprecipitates were dissolved for subsequent immunoblot and electrophoresis assays.

### Quantitative real-time (RT)-PCR

TRIzol reagent (Invitrogen, Carlsbad, CA, USA) and reverse transcribed were employed for extracting total RNA from cultured cells. Real-time PCR analysis was performed on LightCycler 480 II (Roche, Pleasanton, USA) equipped with SYBR green detector (TaKaRa Bio, USA) ^[13]^. The normalization was achieved with GAPDH. All the samples were analyzed in triplicate and data were investigated using the ΔΔCt method. The sequences for all primers are given in Table S2 (Supplementary File).

### The shRNA analysis

Different shRNAs including Negative Control shRNA(shNC), shNF-κB (p50), shWTAP and shNLRP3 were procured from Ribo Targets (Guangzhou, China). For all experimental processes, the control samples were processed using a non-targeting control sequence at the same concentration. The shRNA knock-down efficiency was evaluated using the western blotting technique^[14]^. Table S1 (Supplementary File) of the supplementary file depicts the sequences of shNC, shWTAP, shNF-κB(p50), and shNLRP3.

### Detection of caspase-1 activity

Using commercial assay kits for caspase-1 activity (Beyotime, Shanghai, China), the Caspase-1 activity was assessed following the specifications of the manufacturer. Supernatants from the collected lysed cells were used to assess caspase-1 activity at a wavelength of 450 nm^[15]^.

### LDH assay

LDH release can be used for assessing cell pyroptosis. Therefore, a commercial LDH-cytotoxicity kit (Nanjing Jiancheng Biology Engineering Institute, Nanjing, Jiangsu, China) was used for the quantification of LDH release.

### Enzyme-linked immunosorbent assay (ELISA)

The concentrations of ICAM-1, VCAM-1, MCP-1, and E-selectin in human serum samples and EC culture supernatants were measured by ELISA using Abcam kits.

### Adenovirus-associated virus construction

The adenoviral vector expressing WTAP and the negative control were synthesized by GenePharma (Shanghai Gen-ePharma Co., Ltd., Shanghai, China). AAV Serotype-9 vectors expressing wild-type WTAP and its dominant missense mutation were produced according to the protocol described previously ^[16]^. Mice were injected with 3×10^11^ genome vectors all at once via the tail vein.

### GST-pull down assay

His-NLRP3 protein was purified from *E. coli* and incubated with 10 μg of purified GST or GST-NF-κB protein^[17]^. The GST-associated proteins were then purified using glutathione sepharose 4B, and the bound NLRP3 was detected by western blotting.

### Immunofluorescence assay

The cells’ culture medium was replaced with a fresh medium, and they were incubated for 24 hours before being photographed using a confocal laser scanning microscope (LSM 880 with Airyscan; Zeiss, CA, USA). The nuclei were stained with DAPI for 15 minutes before being imaged with a laser scanning confocal microscope^[18]^ (FV300; Olympus).

### m^6^A level analysis

Total RNA was extracted from endothelial cells with the TRIzol reagent and Poly(A)+ RNA was purified using the GenEl-ute mRNA Miniprep Kit (Sigma, Louis, MO, USA; MRN10). The m6A concentration was assessed using the m6A RNA Methylation Kit (Abcam, ab185912). Specifically, 80 μL of binding solution and 200 ng of sample RNA were introduced into the designated wells, followed by an incubation at 37 °C for 90 minutes to facilitate RNA binding. All wells were washed with wash buffer three times. Next, 50 µl of diluted capture antibody was added to each well and incubated for 60 minutes at room temperature. Subsequently, the detection antibody and Enhancer solution were introduced into each well and allowed to incubate for 30 minutes at room temperature. Following this, the wells were subjected to developer solution in the dark for 1–10 minutes at 25 °C. The reaction was terminated using a stop solution within 2-10 minutes. Finally, the absorbance was measured at 450 nm using a microplate reader.

### Clinical sample collection

In this study, we included 211 patients admitted to our hospital from June 2019 to February 2021 due to chest pain or chest tightness as the main complaint and requiring coronary angiography to assess coronary artery stenosis after completing a series of clinical examinations. The patients were categorized into two groups: the coronary artery disease group (n=143) and the control group (n=68), based on the angiography results. The inclusion criteria for the control group were: no stenosis in the coronary arteries or only myocardial bridging changes. The inclusion criteria for the coronary artery disease group were based on the Judkins criteria: stenosis of at least 50% in at least one of the coronary arteries (left anterior descending, left circumflex, right coronary artery, left main,). The exclusion criteria were: coronary artery bypass grafting and previous coronary angiography; hematologic disorders, neoplasms, severe hepatic and renal insufficiency, active bleeding from any cause, chronic and acute infectious diseases; cardiac arrhythmias; chronic obstructive pulmonary disease, peripheral vascular diseases. This study was approved by the hospital ethics committee, and the included patients signed informed consent forms to participate. Comparison of the general information of the 2 groups, the difference was not statistically significant (*P*>0.05) and was comparable

### Gensini score

The Gensini score is derived from the sum of scores assigned to individual segments. These scores are computed by multiplying a severity score with a segment weighting factor. The segment weighting factors range from 0.5 to 5.0. Severity scores are based on specific percentage reductions in lumen diameter in coronary segments and are allocated as follows: 1 for 25%, 2 for 50%, 4 for 75%, 8 for 90%, 16 for 99%, and 32 for 100%. Consequently, segments supplying larger myocardial areas receive greater weight, and the highest-scoring segments often indicate multiple severe proximal lesions. The degree of stenosis in four vessels: the left anterior descending branch, the left circumflex branch, the right coronary artery the left main stem, was quantitatively scored following internationally recognized methods ^[19]^.

### Statistical analysis

SPSS software (version 13.0; SPSS, Chicago, USA) was used for data analysis. Unless otherwise indicated, data were presented as mean ± Standard Deviation (SD) or median (interquartile range). Continuous variables were analyzed with one-way analysis of variance (ANOVA) or Unpaired Student’s *t*-tests when they distributed normally. Statistical significance was described as a two-tailed P value of less than 0.05.

## Results

### WTAP is highly expressed in LPS-induced endothelial cells and in patients with coronary heart disease(CHD)

To evaluate the significance of WTAP in AS, we analyzed its expression in LPS-induced endothelial cells and in serum samples from patients with CHD. Initially, it was observed that serum m6A levels were significantly elevated in patients with coronary artery disease (Fig.1A). In addition, the results revealed that LPS exposure enhanced the m6A level (Fig.1B) in a dose-dependent manner, thus we chose 1000ng/ml as the concentration for subsequent experiments. Interestingly, among the m6A-regulating proteins (METTL3, METTL4, METTL14, METTL16 and WTAP and FTO and YTHDF1, YTHDF2, YTHDF3, ALKBH5, YTHDC1, YTHDC2), the WTAP genes were upregulated in LPS-induced endothelial cells (Fig. 1C). Next, we further validated the effect of WTAP in CHD. Results indicated that WTAP expression also increased with the development of CHD (Fig. 1D).

**Figure 1.**
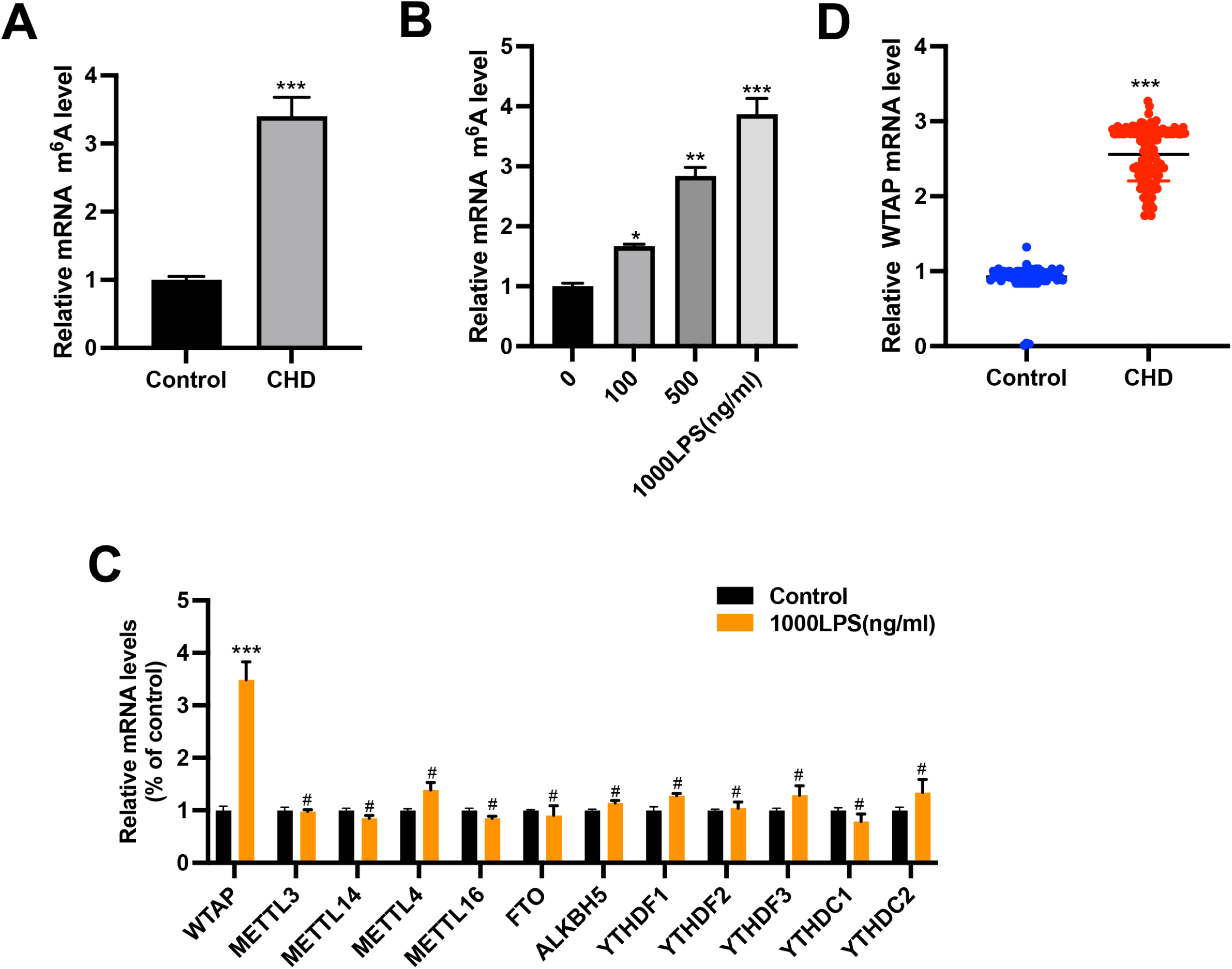

Notably, AS is a chronic inflammatory vascular disease induced by traditional and nontraditional risk factors ^[20]^. Therefore, the expression levels and correlation of inflammation-related factor MCP-1, TNF-α, ICAM-1, VCAM-1, IL-6, IL-18 and WTAP in CHD were explored (Table.1). ELISA results demonstrated that the expression levels of MCP-1, TNF-α, ICAM-1, VCAM-1, IL-6, IL-18 and WTAP were significantly increased in the serum of patients with COPD compared to controls. Furthermore, our findings indicate significant correlations between MCP-1, TNF-α, ICAM-1, VCAM-1, IL-6, IL-18, WTAP, and the Gensini score, highlighting their association with the severity of CHD (Table.2).

**Table.1.** Levels of inflammatory factors.

| Group | TNF-α(pg/ml) | MCP-1(pg/ml) | VCAM-1(pg/ml) |
| --- | --- | --- | --- |
| Control | 145(80.21, 310.67) | 311.09(121.72, 374.32) | 129.04(156.34, 247.83) |
| CHD | 489.2(183.48.2, 497.03) | 549.92(202.16, 584.37) | 329.05(218.52, 498.42) |
| *P* | <0.001 | <0.001 | <0.001 |
| Group | ICAM-1(pg/ml) | IL-6(pg/ml) | IL-18(pg/ml) |
| Control | 100.23(69.38, 280.31) | 169.2(120.56, 321.75) | 149.3(118.5, 327.18) |
| CHD | 330.27(144.29, 376.54) | 378.29(221.9, 532.95) | 431.26(191.4, 545.1) |
| *P* | <0.001 | <0.001 | <0.001 |

**Table.2.**
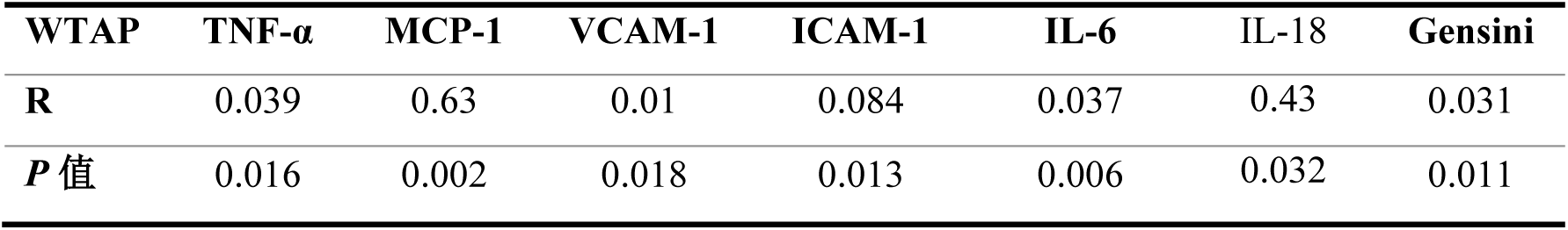
Correlations of WTAP with TNF-α, MCP-1, VCAM-1, ICAM-1, IL-6, IL-18 and gensini scores.

| WTAP | TNF-α | MCP-1 | VCAM-1 | ICAM-1 | IL-6 | IL-18 | Gensini |
| --- | --- | --- | --- | --- | --- | --- | --- |
| *R* | 0.039 | 0.63 | 0.01 | 0.084 | 0.037 | 0.43 | 0.031 |
| *P* 值 | 0.016 | 0.002 | 0.018 | 0.013 | 0.006 | 0.032 | 0.011 |

### Lipopolysaccharide triggers pyroptosis and inflammatory response in endothelial cells

Next we further explored the mechanism of inflammatory response in AS. Pyroptosis is pro-inflammatory-regulated cell death and is believed to be involved in AS^[21]^. Thus, the relationship between LPS, the inflammatory response, and pyroptosis in Endothelial cells was investigated. Our results suggested that LPS exposure enhanced the inflammatory response as evidenced by the increased production and mRNA levels of vascular cell adhesion molecule (VCAM)-1, intercellular adhesion molecule (ICAM)-1, monocyte chemoattractant protein (MCP)-1, and E-selectin (Fig. 2A–B) in a dose-dependent manner, thus confirmed a role of LPS induced inflammatory responses in Endothelial cells. Next, we further evaluated the LPS effects on cell pyroptosis via determining the expression level of the pyroptosis-related gene. The western blot findings demonstrated that LPS enhanced the levels of several pyroptosis-related proteins [*NLRP3, cleaved caspase-1, p20, p17, IL-18, C-terminal GSDMD (GSDMD-C), cleaved IL-1β,] and full-length GSDMD*) upon Endothelial cells exposure to LPS. Furthermore, there were no major shifts in the expression levels of inactive IL-1β and caspase-1 precursors. In addition, the expression of genes caspase-1, NLRP3, GSDMD, IL-18, and IL-1β increased dose-dependently at the protein and mRNA level (Fig.2C-D). Studies have previously shown that such pyroptosis was complemented by plasma membrane pore formation, which caused the release of intracellular contents, e.g. lactate dehydrogenase (LDH), and that LDH activity was dose-dependently increased by LPS (Fig. 2E). Furthermore, the activity of caspase-1 was significantly increased by LPS (Fig. 2F). The data showed that LPS promoted cell pyroptosis and inflammation, and our findings suggested that LPS-induced cell pyroptosis may play a role in the inflammatory response seen in Endothelial cells.

**Figure 2.**
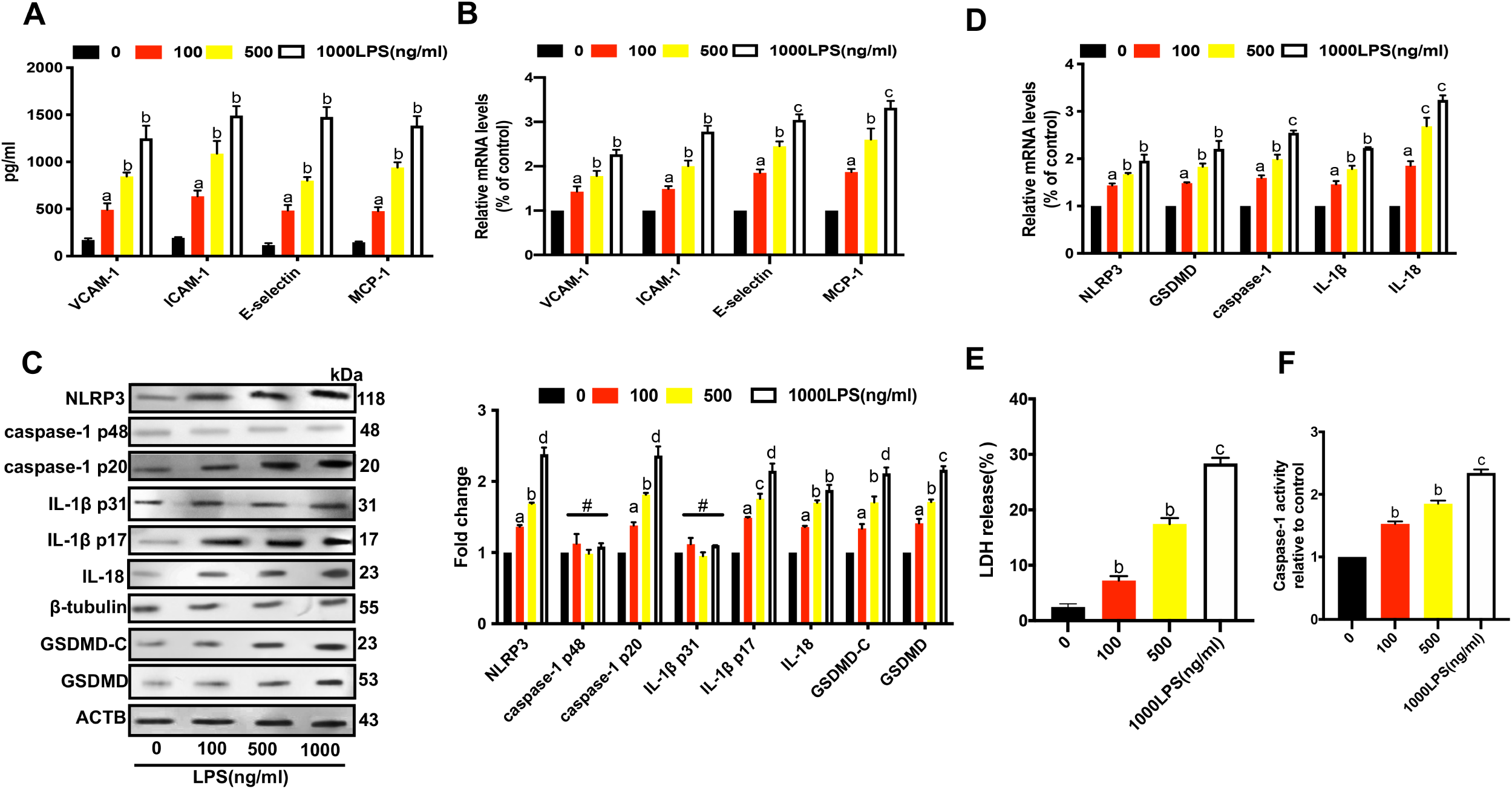

### WTAP aggravates LPS-induced pyroptosis and inflammation in endothelial cells

To further understand WTAP’s functional role and potential mechanisms of action in endothelial cells, we first investigated its role in LPS-induced pyroptosis and inflammation. The knockdown of WTAP (shWTAP) downregulated the expression of WTAP (Supplementary Figure 1A). As shown in Fig.3A-B, LPS significantly enhanced the expression of WTAP in a dose-dependent manner. When Endothelial cells were treated with both LPS and shWTAP, the production of inflammatory cytokines (VCAM-1, ICAM-1, E-selectin and MCP-1) were significantly reduced as compared to the LPS group alone (Fig.3C). Importantly, our qRT-PCR results matched those of ELISA (Fig.3D), implying that shWTAP inhibited LPS-induced inflammation in Endothelial cells. Next, we further assessed the WTAP effects on pyroptosis induced by LPS. Both LPS and shWTAP treatment resulted in a downregulation of pyroptosis-related genes (*caspase-1, NLRP3, IL-1β, GSDMD,* and *IL-18*) at the protein level, with lower levels than in the LPS group (Fig.3E). These results were consistent with qRT-PCR (Fig.3F). Furthermore, LPS-induced LDH release and caspase-1 activity were both decreased by shWTAP(Fig.G-H). These results suggested that WTAP may be involved in pyroptosis and inflammation induced by LPS in Endothelial cells.

**Figure 3.**
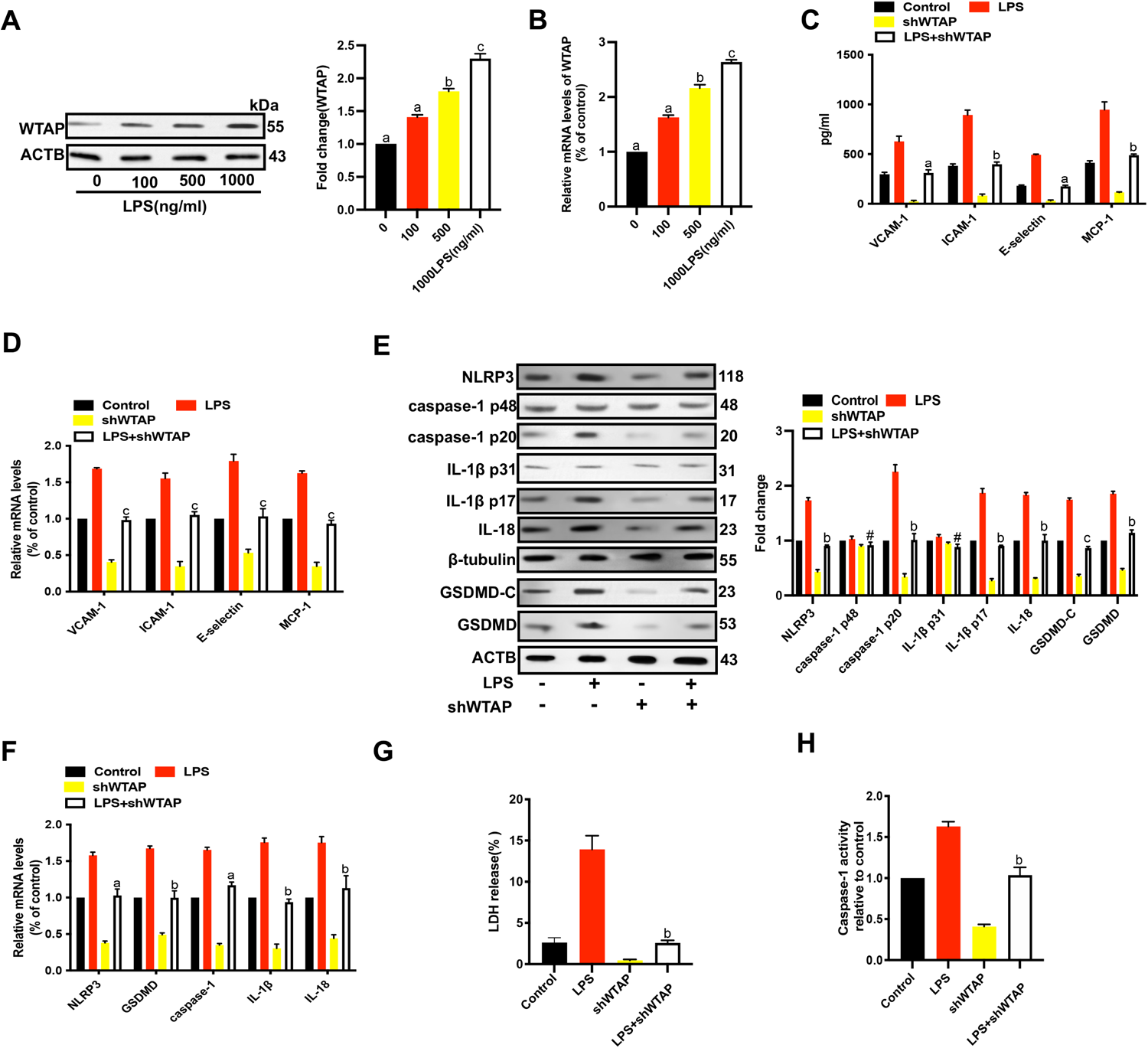

### LPS-induced pyroptosis and inflammation in endothelial cells is regulated by WTAP via NF-κB

The previous experiments have suggested that NF-KB signaling pathway plays an important role in the formation of AS by activating the production of pro-inflammatory factors^[22]^. Next, we investigated the involvement of NF-κB in pyroptosis and inflammation and found that LPS increased the protein and mRNA levels of NF-κB p50 in Endothelial cells in a dose-dependent manner (Fig.4A-B), implying that LPS increased the potential activity of NF-κB p50 in Endothelial cells. Thus we further investigated the relationship between WTAP and NF-κB induced inflammatory response.Our results found that shWTAP inhibited the NF-κB p50 activity, indicating that shWTAP plays a role in NF-κB p50 inhibition(Fig. 4C-D). After this, we used small interfering RNAs (shRNAs) to knockdown p50 and confirmed a reduced effect of NF-κB p50 as detected by qRT-PCR (Supplementary Figure 1B). When compared to LPS alone, the production of the inflammatory cytokines (*VCAM-1,ICAM-1, E-selectin, and MCP-1*) were all markedly decreased in Endothelial cells after treatment with LPS and shNF-κB(p50) (Fig.4E). Furthermore, the findings of qRT-PCR results were consistent with ELISA (Fig.4F), indicating that knocking down p50 reduced LPS-induced inflammation. Next, we further assessed the effects of NF-κB p50 on LPS induced pyroptosis. Analysis of pyroptosis-related genes (*NLRP3, IL-1β, IL-18, GSDMD* and *caspase-1*) at the mRNA level revealed that they were downregulated in the LPS group compared with the control. These results were in good agreement with those of western blotting analysis (Fig.4G-H). Furthermore, the results showed that shNF-κB (p50) attenuated the LPS-induced LDH release (Fig. 4I) and that the LPS-induced caspase-1 activity was suppressed by shNF-κB (p50) (Fig.4J), suggesting that LPS-induced pyroptosis and inflammation might be mediated by NF-κB (p50). Based on these findings, we hypothesized that WTAP enhanced pyroptosis and inflammation induced by LPS in endothelial cells via the NF-κB pathway.

**Figure 4.**
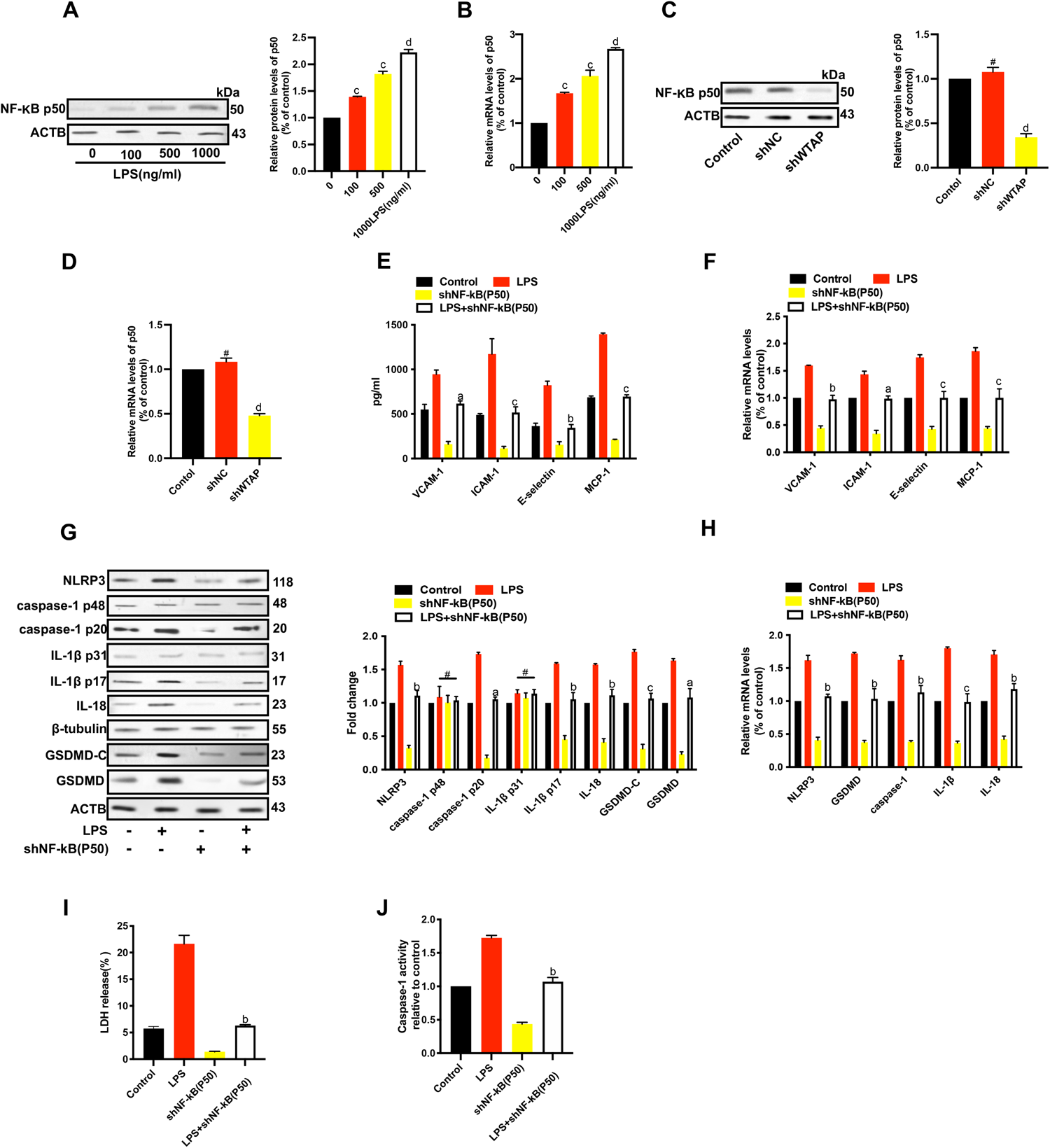

### WTAP enhanced pyroptosis and inflammation in endothelial cells via the NF-κB/NLRP3 signaling pathway

shRNAs were designed to knock down NLRP3 (shNLRP3) to further elucidate the impact of NLRP3 on LPS-induced pyroptosis and inflammation, and this knockdown effect was validated by qRT-PCR (Supplementary Figure 1C). Fig.5A-B shows that shNLRP3 significantly reduced LPS-induced production of VCAM-1, ICAM-1, E-selectin, and MCP-1 when compared to the control group, and this was consistent when employing qRT-PCR. The above data indicated that NLRP3 was involved in the process of LPS-mediated inflammation. Moreover, we also assessed whether there were any regulatory effects of NLRP3 on LPS and subsequently on pyroptosis. As shown in Fig.5C-D, the expressions of genes related to pyroptosis (*caspase-1, IL-1β, IL-18,* and *GSDMD*) were upregulated by LPS, while downregulated by shNLRP3, at both protein and mRNA levels. Furthermore, LPS-induced LDH release was attenuated by shNLRP3 (Fig.5E). Additionally, the activity of caspase-1 induced by LPS was decreased by shNLRP3 in Endothelial cells (Fig.5F). These results demonstrated that LPS-induced pyroptosis and inflammation might be mediated by NLRP3 in Endothelial cells. Furthermore, dual-luciferase assays indicated that the levels of both the 5′UTR and 3′UTR of NLRP3 were affected by WTAP (Fig. 5G-H). WTAP could methylate 3’-UTR of CAV-1 and downregulate CAV-1 expression to activate NF-κB signaling pathway^[23]^. Studies have shown that NF-κB regulates downstream NLRP3 and hence plays a vital role in the processes related to cardiovascular disease ^[24–25]^. Therefore, we further analyzed the precise relationship between NLRP3 and NF-κB in Endothelial cells and our data indicated that p50 knockdown decreased the expression of NLRP3 in these cells (Fig.4G). In addition, a potential physical interaction between NF-κB and NLRP3 was explored using immunofluorescence, co-immunoprecipitation, and GST-pulldown assays. We found that, in Endothelial cells, p50 and NLRP3 were highly co-localized (Fig.5I) and that an anti-p50 antibody could pull down the endogenous NLRP3 protein. These immunoprecipitations were resolved using SDS-PAGE, and the results revealed that NLRP3 was successfully immunoprecipitated (Fig.5J). Furthermore, a glutathione S-transferase (GST) pull-down assay using purified p50 and NLRP3 protein revealed that p50 directly interacted with NLRP3 (Fig.5K), demonstrating that p50 played a substantial regulatory role within the NF-κB/NLRP3 signaling pathway by directly interacting with NLRP3. According to above findings, WTAP may have increased LPS-induced pyroptosis and inflammation in Endothelial cells by modulating the NF-κB/NLRP3 pathway as suggested by these findings.

**Figure 5.**
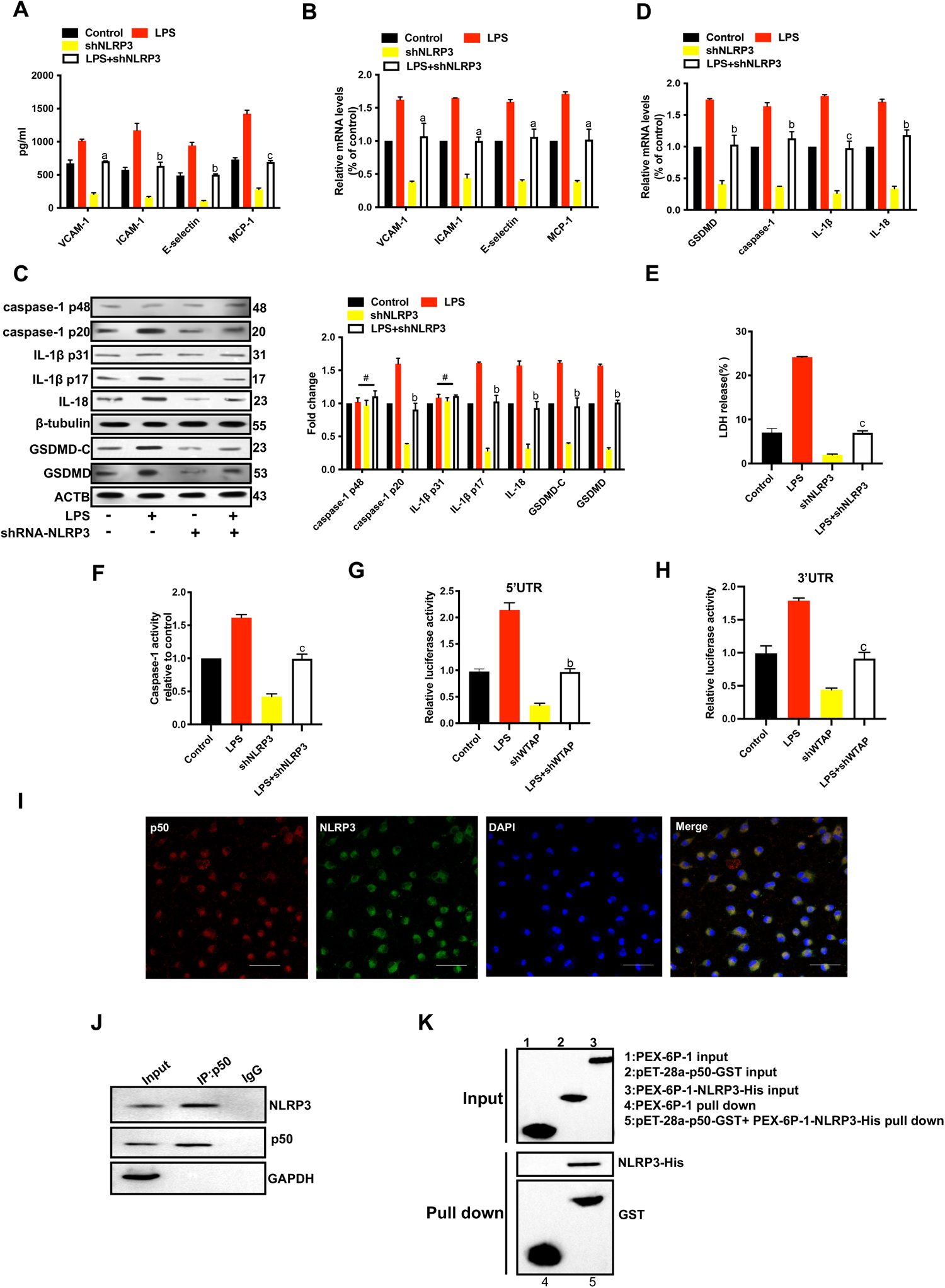

### WTAP augmented atherosclerotic lesions

To determine whether WTAP had an effect on the development of atherosclerosis, we established an in vivo model of AS by feeding ApoE^-/-^ mice a high-fat diet to observe the effect of WTAP on atherosclerotic plaques. We established an WTAP-interference model by tail vein injection of WTAP adenovirus interference vector. The size of the aortic plaque was evaluated by hematoxylin-eosin staining (HE) and the collagen fibre content was measured by Masson staining. HE stains revealed a significant decrease in damage at the aortic valve after administration of AAV-si-WTAP-treated compared to HFD+AAV-Mock group. More importantly, AAV-si-WTAP injected mice were to have smaller atherosclerotic lesions in the aortic valve than HFD+AAV-Mock group (Fig. 6A). Masson staining showed that the collagen fibre content in the AAV-si-WTAP treated mice was significantly lower than that in the HFD+AAV-Mock (Fig.6B). As expected, immunohistochemical analyses of NLRP3 and P50 showed that in the AAV-si-WTAP-treated group the level of NLRP3 and P50 was decreased (Fig. 6C) in AAV-si-WTAP mice.

**Figure 6.**
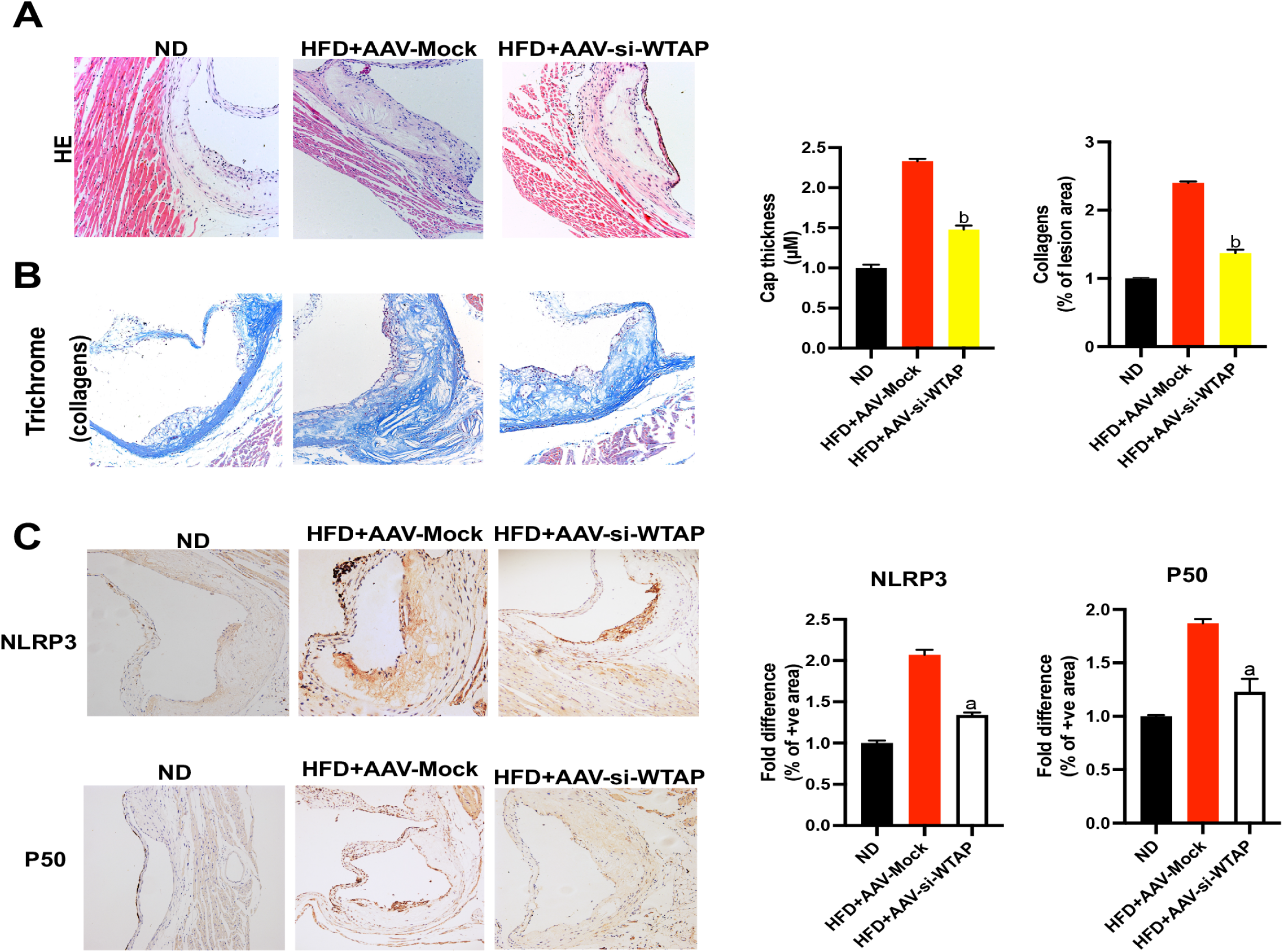

## Discussion

Atherosclerosis represents a lipid-induced inflammatory disease where the balance between pro-inflammatory and anti-inflammatory mechanisms is critical for the development of clinical events ^[26]^. Pyroptosis is a lytic programmed cell death that causes swelling of cells, leading to the release of inflammation-related cytokines such as IL-18 and IL-1β ^[27]^, and the suppression of pyroptosis by moderate hypothermia reduces ischemic injury ^[28]^. In acute myocardial infarction, Sirtuin (SIRT)6 attenuate (TREM)-1-mediated pyroptosis and EC inflammation ^[29]^. Aloin, a plant extract, has been found to inhibit LPS-induced inflammation through the suppression of JAK1-STAT1/3 activation^[30]^ and postnatal inflammation following intrauterine inflammation exacerbated the development of AC in ApoE ^-/-^ mice ^[31]^. In addition, LPS-induced inflammation affected cholesterol accumulation and decreased the levels of the adenosine triphosphate (ATP)-binding cassette transporter A1 (ABCA1) in AS ^[32]^. The results of our current study confirmed the findings of a previous study, showing that LPS increased inflammatory responses during the pyroptosis process in a dose-dependent manner. Caspase-1, a member of the caspase protease family and an “inflammatory caspase” with the CARD domain in the region of NH_2_-terminus, was also upregulated. This caspase is an important part of the caspase-1 inflammasome, which regulates pyroptosis through the canonical inflammasome pathway. Moreover, caspases-1 plays an important role in the activation of Endothelial cells, and its overexpression promotes the development of AS ^[33]^. In this study, we observed that the activity and expression of caspase-1 were increased by LPS and this caspase cleaved pro-inflammatory cytokines, which led to the activation of IL-18 and IL-1β, inducing an inflammatory response^[34]^. Our findings also revealed that LPS induced the cleaved forms of IL-18 and IL-1 (P31) in a dose-dependent manner. Previous studies have revealed that pyroptosis was triggered by both the canonical and noncanonical inflammasome pathways, but caspase-1 exclusively triggered the canonical pathway. The CARD domain removal caused by NLRP3 activation leads to autocleavage of caspase-1, resulting in the formation of an activated caspase-1 p10/p20 tetramer, which is responsible for pyroptosis. This, in turn, cleaves GSDMD, producing N-terminal p30 and C-terminal p20 fragments, where the N-terminal p30 fragment makes functional pores in membranes leading to cell pyroptosis ^[35]^. Recent studies, including our own have found that LPS induces pyroptosis through the canonical inflammasome pathway and that this LPS-induced response might be due to cell pyroptosis activation, suggesting that LPS-induced pyroptosis could be the responsible cellular mechanism for inflammatory response.

The NLRP3 inflammasome is a multi-protein complex that triggers inflammatory responses and can be activated by the canonical pathway after caspase-1 activation, resulting in pyroptosis ^[36]^. Myocardial ischemic injury is exacerbated by the activation of NLRP3-inflammasome-regulated pyroptosis ^[37]^. Furthermore, the STING-IRF3 by activation of NLRP3 contributes to LPS-induced cardiac inflammation, dysfunction, and pyroptosis ^[38]^. Our current research has revealed a close association of the NLRP3 inflammasome with cell pyroptosis, and thus played a critical contribution in the development of cardiovascular disease. Previous research studies have reported that the NLRP3 inflammasome signaling pathway components were mainly localized to unstable atherosclerotic plaques of human carotid ^[39]^. Our results further demonstrated that LPS-induced pyroptosis and inflammation might be mediated by NLRP3 in Endothelial cells. Therefore, we speculate that LPS regulates EC pyroptosis via NLRP3 upregulation, which then activates a series of signaling cascades, including the pyroptosis-related inflammasome pathway through caspase-1 upregulation, which promotes inflammation and accelerates the progression of AS. However, the exact underlying mechanisms stand obscure.

Various chronic inflammatory diseases can trigger the development and progression of AS ^[40–41]^. Currently, the specific mechanism linking inflammatory response to the occurrence of AS has not been fully clarified. Research has shown that m6A may have clinical benefits in the management of AS. WTAP regulates m6A modification expression in THP-1 macrophages ^[42]^ to exert anti-inflammatory effect of APS. WTAP mediated the injurious effects of myocardial I/R by augmenting cell apoptosis and ER stress through a mechanisms involving the modulation of m6A modification on ATF4 mRNA ^[43]^. Wu et al. confirmed that m6A methylation levels were significantly decreased in leukocytes from patients with AS and in mice ^[44]^.

In this study, we found that WTAP enhanced inflammatory responses and pyroptosis induced by LPS via NLRP3. These results provided evidence that shWTAP has a protective role against the inflammatory process of AS and in addition, inhibition of NF-κB reduced this NLRP3-mediated pyroptosis ^[45]^. Melatonin also reduces the inflammasome-induced pyroptosis by blocking NF-κB ^[46]^. Consistent with the above study, we found that p50 knockdown remarkably alleviated NLRP3-mediated pyroptosis and inflammation induced by LPS, suggesting that NLRP3 might be regulated by NF-κB (p50). To support this, numerous studies have shown that inhibition of the NF-κB/NLRP3 pathway attenuates myocardial ischemia-reperfusion injury in mice^[47]^, whereas sinigrin can suppress macrophage-associated inflammatory responses by inhibiting the NF-κB/NLRP3 pathway ^[48]^. In this study, we observed a reduction in NF-κB p50 activity due to shWTAP. Crucially, shWTAP mitigated NLRP3-mediated pyroptosis and inflammation via NF-κB in endothelial cells. Taken together, our findings highlight the involvement of WTAP in LPS-induced pyroptosis and inflammation through the NF-κB/NLRP3 pathway in endothelial cells.

## Conclusion

In this study, we identified for the first time that WTAP significantly increased pyroptosis and inflammation induced by LPS. In addition, NF-κB p50 activity was inhibited by shWTAP, and p50 interacted directly with NLRP3 in Endothelial cells. In conclusion, WTAP can play a key role in EC pyroptosis and inflammation through the NF-κB/NLRP3 signaling pathway.

## Ethics statement

This study was conducted with approval of the academic ethics committee of Henan Provincial People’s Hospital. All procedures strictly followed the code of Declaration of Helsinki.

## Consent to publish

Not applicable.

## Data Availability Statement

Not applicable

## Author Contributions

Fengxia Guo and Mei He conceived and supervised the study; Fengxia Guo designed experiments; Fengxia Guo and Yanhua Sha performed experiments; Fengxia Guo and Bing Hu analyzed the data; Fengxia Guo wrote the manuscript; and Fengxia Guo and Yanhua Sha revised the manuscript. All authors read and approved the final manuscript.

## Acknowledgements

Not Applicable.

